## Supplemental Figure 1-5 for "Tracking inflammation resolution signatures in lungs after SARS-CoV-2 omicron BA.1 infection of K18-hACE2 mice"

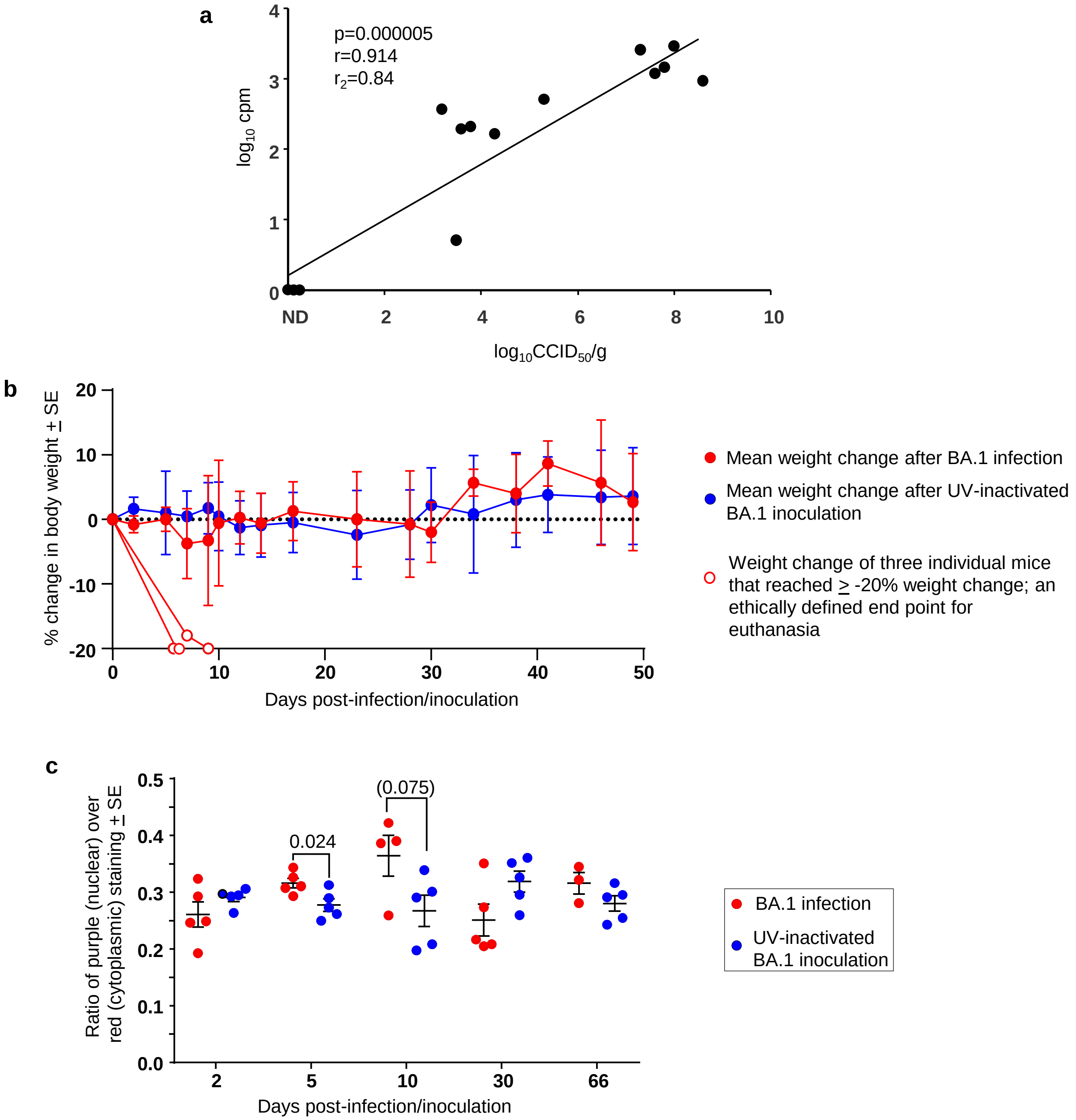


**S1 Fig.** **a** Correlation of viral loads measured by RNA-Seq (counts per million reads) (using one lung lobe) vs. viral titers by $CCID_{50}$ assays (using another viral lobe) for each mouse for 2, 5, and 10 dpi. Despite different assay methods and different lung lobes being used, correlation was high. Statistics by Pearson correlation. If 30 (n=5) and 66 (n=3) dpi data are added (all very low or zero for both titer and cpm), then r=0.953, p = 7.194E-12. **b** Data from Stewart et al. 2023 Supplementary Fig. 1a, regraphed to show means for BA.1-infected and UV-inactivated BA.1 inoculated mice. Three out of 25 infected mice in this cohort were euthanized due to reaching >20% weight loss, which was associated with brain infection; these 3 mice are plotted individually (open circles) and such mice were excluded from the current study. For UV-inactivated BA.1 inoculated mice (n=24). For each time point 3-12 mice were weighed to produce a mean and SE. **c** The ratio of purple (nuclear) to red (cytoplasmic) staining in H&E stained sections provides a crude automated (Aperio Positive Pixel Count) measure of leukocytes infiltrates as leukocytes have a high nuclear to cytoplasmic area ratio (Dumenil et al., 2023). Statistics by t tests.

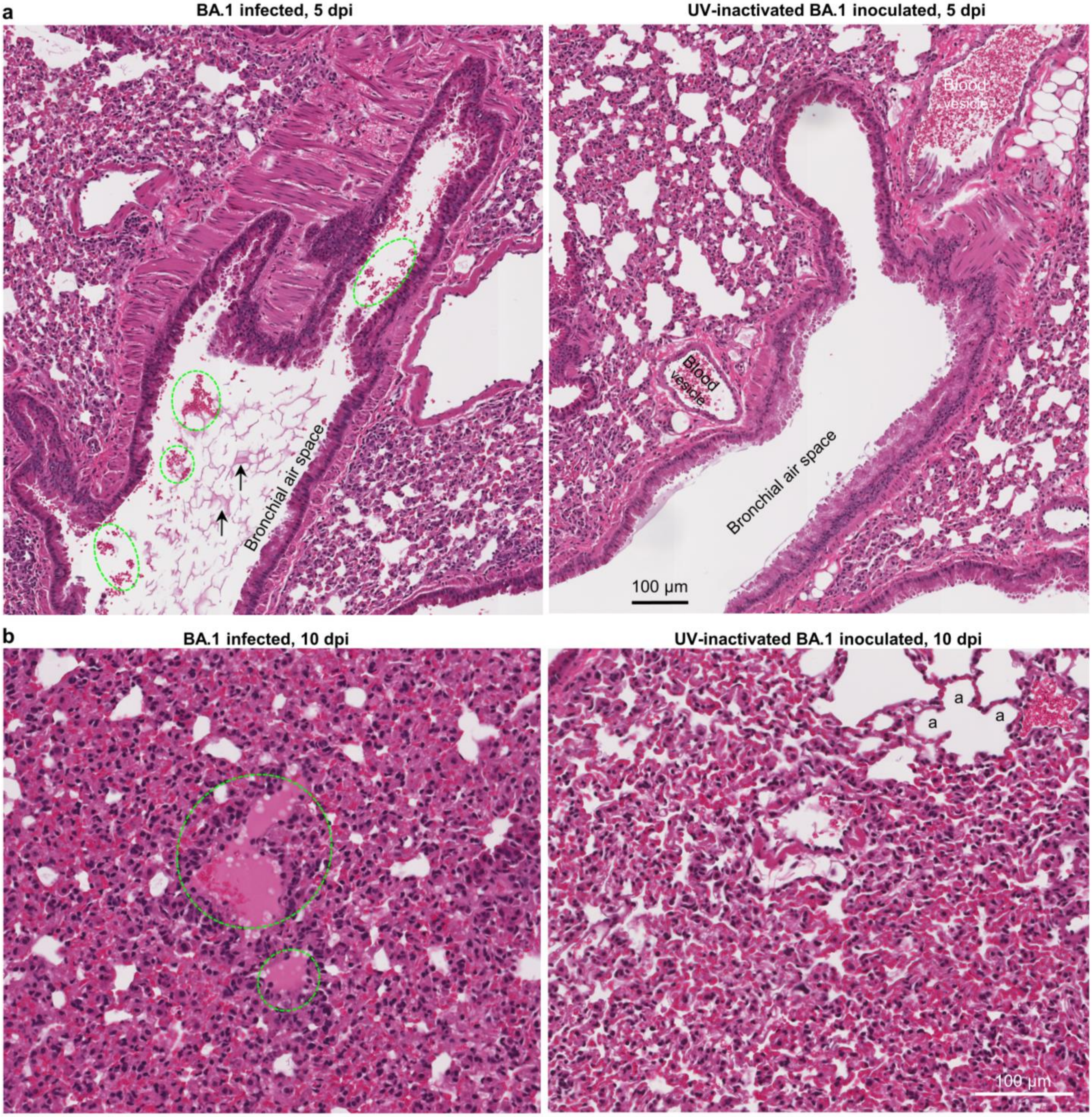


**S2 Fig.** **a** BA.1 infected mice occasionally (2/5 mice) show some red blood cells (dashed green ovals) and serum (arrows) in bronchi; not seen in mice inoculated with UV-inactivated BA.1. **b** BA.1 infected mice show occasionally alveolar oedema (dashed green ovals) in 4/4 mice at 10 dpi, a feature rarely observed in mice inoculated with UV-inactivated BA.1. Left hand image also shows an area of severe lung consolidation, with loss of alveolar air spaces. a – alveolar air sacks.

**a** PBS inoculated day 5

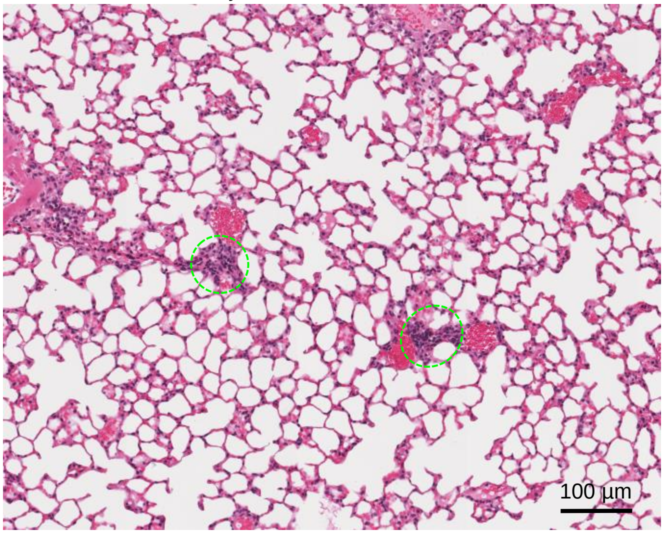


**b** PBS inoculated day 5

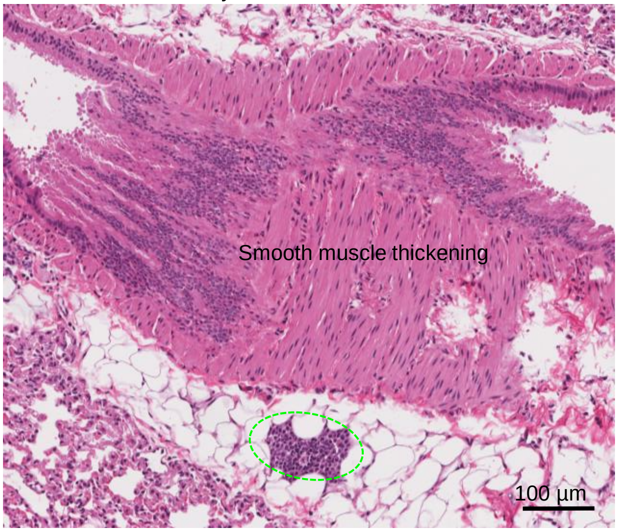


**c** PBS inoculated day 5

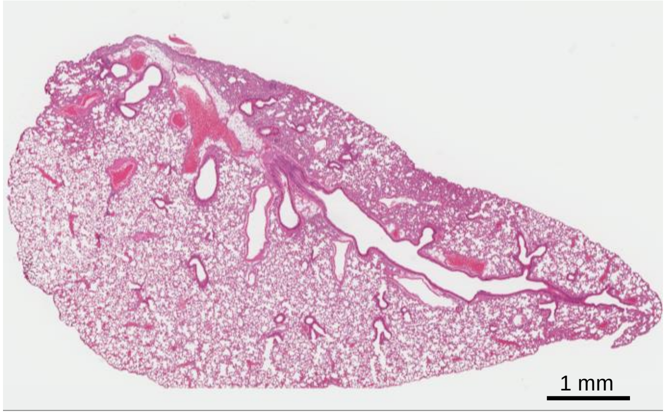


**d** Naïve mouse (5 months old)

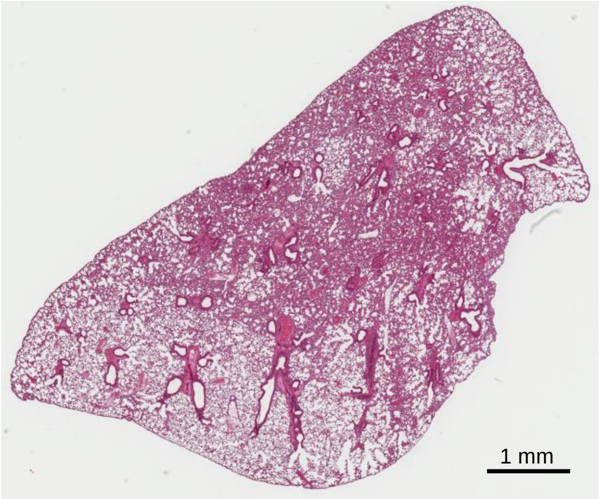


**e** Naïve mouse (5 months old)

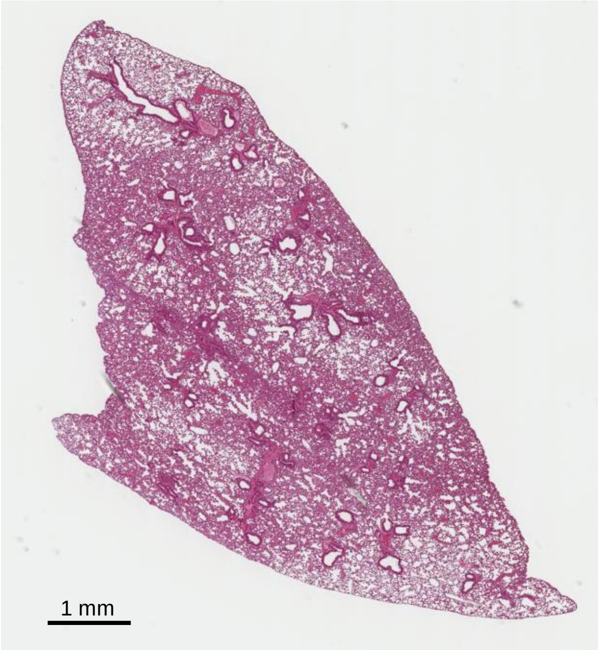


**S3 Fig.** H&E staining of control lungs from K18-hACE2 mice**. a, b.** Small foci of cellular infiltrates can occasionally occur in mice inoculated with PBS (dashed green ovals). **c** Lung consolidation is generally not observed after inoculation of PBS. **d,e** Age-related loss of white space.

**a**

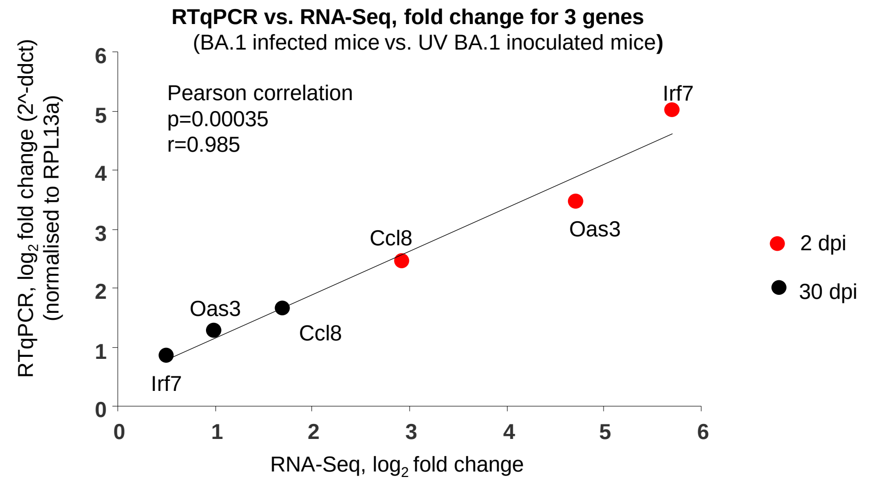


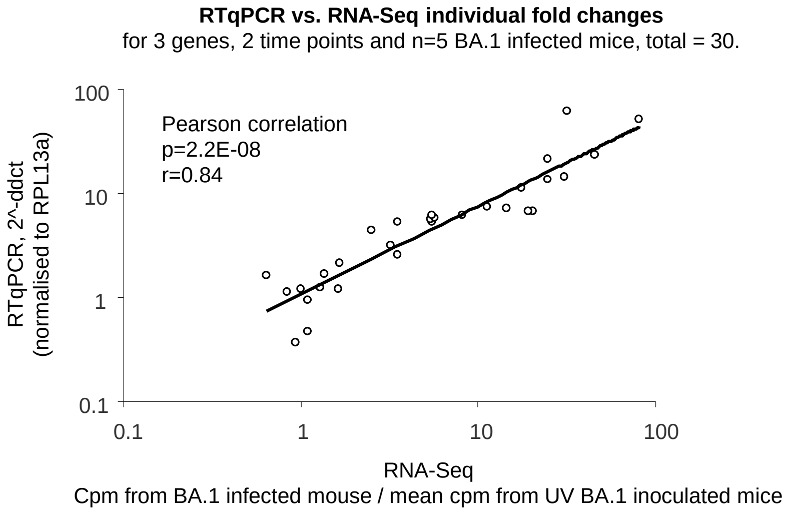


**b**

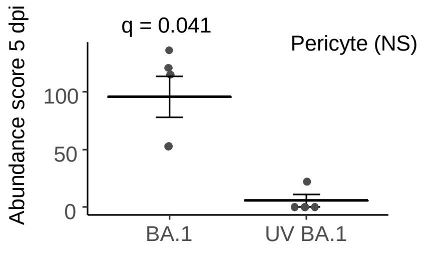


**S4 Fig.** **a** RT qPCR validation of RNA-Seq data using 3 genes and 2 time points. Top graph correlation between fold change data generated using the two methods; 3 genes and 2 time points, mean from n=4/5 mice per group. Bottom graph individual fold change data for each BA.1 infected mouse. Data for UV BA.1 inoculated mice was meaned for each gene and time point. **b** Cellular deconvolution (SpacialDecon) as for Fig. 3c also identified significant increased expression of gene (mRNA) signatures associated with pericytes.

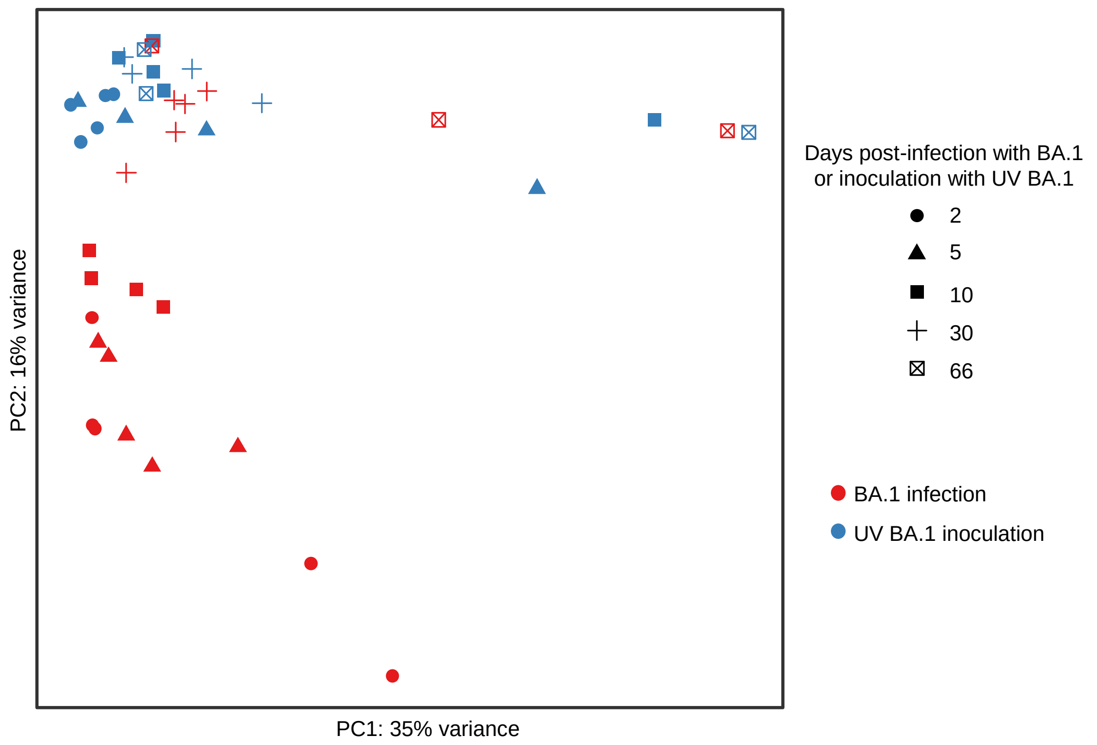


**S5 Fig**. First two principle components of normalised VST-transformed counts from all experimental groups. No clear segregation for 66 dpi with BA.1 (⊠) and 66 days after inoculation with UV-inactivated BA.1 (⊠).
